# Towards understanding of NK cell antigenic specificity

**DOI:** 10.64898/2026.05.29.728791

**Authors:** Maria O. Ustiuzhanina, Irina A. Shagina, Evgeniy Nikitin, Evgeniy Klimuk, Maria Salnikova, Olga V. Britanova, Yendry Ventura-Carmenate, Elena I. Kovalenko, Dmitriy M. Chudakov

**Affiliations:** Center for Molecular and Cellular Biology, Moscow, Russia; Shemyakin-Ovchinnikov Institute of Bioorganic Chemistry, Russian Academy of Sciences, Moscow 117997, Russia; Institute of Translational Medicine, Pirogov Russian National Research Medical University, Moscow 117513, Russia; Biotech Campus LLC, Moscow, Russia; Department of Immunology, Faculty of Biology, Lomonosov Moscow State University, Moscow 119991, Russia; Abu Dhabi Stem Cell Center, Al Muntazah, United Arab Emirates

**Keywords:** NK cells, KIR, scRNA-Seq, transcriptome, adaptive NK cells, memory NK cells, in silico modeling

## Abstract

NK cells can form clonal populations demonstrating features of adaptive immunity, including long-term memory and at least partial antigenic specificity. Given the limited individual diversity of activating receptors, the nature of NK cell antigenic specificity remains elusive. To explore this riddle, we combined scRNA-Seq of *ex vivo* FACS-sorted NK cell subsets expressing specific KIR receptors, single-cell cloning and bulk RNA-Seq of *in vitro* cultured KIR2DS4⁺ NK cell clones, transcriptomic profiling of antigen-stimulated NK cells, and *in silico* modeling of glycosylated KIR2DS4-peptide-HLA complexes. scRNA-Seq resolved 12-15 clusters per KIR⁺ subset with highly heterogeneous KIR, KLRC and NCR expression patterns, consistent with clonal lineages. Notably, those clusters demonstrated over 30 differentially expressed glycosyltransferase genes, potentially involved in post-translational modification of NK cell receptors. Single-cell-derived KIR2DS4⁺ cultures exhibited clone-specific cytotoxic, chemokine and KIR receptor genes, and transcriptional differences in > 40 glycosyltransferases. In peptide culturing autologous assays, SARS-CoV-2 (KTFPPTEPK) and EBV (CRAKFKHLL) peptides elicited NK cell proliferation and distinct transcriptional programs linking cytotoxicity genes, KIR2DS4 and glycosyltransferases. Structural modeling revealed that N-linked glycosyl residues in specific regions of KIR2DS4 may alter its contacts and interaction with MHCI and the presented peptide. We conclude that KIR⁺ human NK cells comprise clonally imprinted populations with distinct glycosyltransferase expression profiles, and site-specific KIR2DS4 glycosylation may modulate interaction with peptide-MHCI complexes, suggesting a post-translational layer of clonal NK cell diversification as a clue to their antigenic specificity.

## Introduction

Natural killer (NK) cells are cytotoxic lymphocytes that provide rapid protection against viral infection and malignant transformation^1–3^. Their activity is controlled by the integration of signals from an array of activating and inhibitory receptors, including killer cell immunoglobulin-like receptors (KIRs) and CD94/NKG2 heterodimers, which interact with host major histocompatibility complex (MHC) class I molecules. This receptor architecture positions NK cells at the interface between innate and adaptive immunity, enabling them to respond quickly while still encoding some features of immune memory.

NK cells can undergo clonal expansion, long-term persistence and “recall” responses after viral infection^4,5^. In murine cytomegalovirus (mCMV) infection, Ly49H⁺ NK cells pass through expansion, contraction and memory phases that closely resemble CD8⁺ T cell responses^6,7^. Using retroviral barcoding of individual Ly49H^+^ NK cells, clonal expansion was determined following mCMV infection^8^. Applying barcoding of stem cells in rhesus macaques it was demonstrated that rhCMV infection drives the emergence and long-term maintenance of expanded NK clones, rather than a transient, polyclonal burst^9^. Those cells can be further cultivated^10^. In humans, infection with human cytomegalovirus (hCMV) drives the accumulation of NKG2C⁺ CD57⁺ “adaptive” NK cells with a stable epigenetic and transcriptional imprint and altered signaling module composition^11–16^. Lineage-tracing approaches using mitochondrial DNA mutations have further shown that many NK cell subsets in adult humans represent long-lived clonal lineages, indicating that NK cells in peripheral blood are clonally diverse^17^.

A central determinant of NK cell specificity is the repertoire of KIRs and their interaction with peptide-MHC class I (pMHC) complexes. KIR genes are highly polymorphic in population and are inherited in A and B haplotypes, giving rise to extensive inter-individual variation in the number and type of activating and inhibitory KIRs^18–20^. At the cellular level, stochastically chosen expression of KIRs, NKG2 and other receptors, generates a vast combinatorial repertoire of NK cell clones, each with a distinct set of self-MHC contacts and activation thresholds^21–24^. Structural and functional studies have shown that in some cases KIRs can discriminate between different peptides presented by the same HLA allotype, indicating that NK cell recognition can reach certain level of peptide specificity^25–34^.

Most of our mechanistic understanding of clonal NK cell memory, however, comes from models centered on a single receptor-ligand axis: Ly49H-m157 in mice or NKG2C-HLA-E in humans. In contrast, clonal behavior and antigen responsiveness of NK cells bearing activating MHC-class-I recognizing KIRs, such as KIR2DS4, remain much less explored, despite genetic evidence that these receptors contribute to hCMV-driven imprinting of the human KIR repertoire^35^. Whether human NK cell clones expressing activating KIRs may also form stable, functionally distinct lineages analogous to NKG2C⁺ adaptive NK cells^17^ is an open question. In particular, KIR2DS4 is an attractive model receptor: it can recognize subsets of HLA-C and possibly HLA-A*11 molecules in a peptide-dependent manner^34^. Yet, the clonal architecture of KIR2DS4⁺ NK cells, and the transcriptional programs that accompany their antigen-driven expansion have not been systematically characterized.

Beyond epigenetic reprogramming, post-translational modifications of NK cell receptors represent an underexplored mechanism of clonal diversification. We have extensively summarized the NK cell antigen-specific function and proposed the influence of post-translational modifications on NK cell receptor diversity^24^. NK receptors could be N-and O-glycosylated, and glycan composition can affect receptor folding, surface stability and ligand binding^36–39^. For KIR3DL1, allele-specific differences in N-linked glycosylation have been shown to modulate binding to HLA-Bw4 ligands and to tune inhibitory signaling strength^40^. These observations raise the possibility that clonal NK cell responses are epigenetically encoded not only in respect of expression of particular patterns of inhibitory and activating receptors, but also in respect of clone-specific glycosylation programs fine-tuning the pMHC recognition.

Here, we address these gaps by combining single-cell RNA sequencing of sorted KIR2DS4⁺ and KIR2DL1⁺ NK cell subsets, single-cell cloning of KIR2DS4⁺ NK cells, and autologous antigen-specific proliferation assays with viral peptides presented in defined MHCI contexts. This allowed us to focus on the clonal structure of activating KIR⁺ NK cells, to link antigen-specific proliferative responses to distinct transcriptional programs, and to map clone-specific patterns of glycosyltransferase expression. Finally, using structural modeling of KIR2DS4-peptide-HLA complexes with defined glycosylation patterns, we explored how site-specific receptor glycosylation might modulate KIR2DS4-peptide interaction. Together, these data provide a first framework for understanding how clonal diversity, antigen experience and post-translational modification may intersect to shape KIR-mediated NK cell specificity.

## Results

### 1. “Zoom in” scRNA-Seq of sorted KIR2DS4^+^ and KIR2DL1^+^ NK cells reveals clonal receptor patterns

Emerging evidence suggests that, upon expansion in response to specific stimuli, NK cells can form long-lived, functionally and phenotypically distinct clones^8,17,35,41^. Models used to investigate such clonality have primarily focused on response to CMV infection, in which NKG2C⁺ clonal NK cells accumulate^8^. At the same time, clonal NK cells that would express other activating HLA-recognizing receptors, such as KIR2DS or KIR3DS^35^ remain less explored. One of the reasons for this is the huge diversity of NK cell clones in a given individual^42,43^, and correspondingly low frequency of each NK cell clone.

To overcome this limitation, we performed scRNA-Seq for the small FACS-sorted subsets of activating KIR⁺ NK cells, in order to “zoom in” the landscape of their clonal diversity. Peripheral blood NK cells were isolated from two donors with documented KIR2DS4 expression and distinct HLA genotypes. Donor 1 carries HLA-A*11, a known ligand for KIR3DL2 and a putative candidate for interaction with KIR2DS4^44,45^, whereas Donor 2 carries the HLA-C2 allele, a canonical ligand for KIR2DL1 with comparatively lower affinity for KIR2DS1 and KIR2DS4^46,47^.

From Donor 1, we sorted KIR2DS4^+^NKG2C^−^ and KIR2DS4^+^NKG2C^+^ NK cells (KIR3DL2^−^CD3^−^CD14^−^CD20^−^CD56^+^), while from Donor 2 we sorted KIR2DS4^+^KIR2DS1^−^KIR2DL1^−^NKG2C^−^ and KIR2DS4^−^KIR2DL1C^245+^ NK cells (CD3^−^CD14^−^CD20^−^CD56^+^) (**Fig. 1a, Supplementary Fig. 1a, Supplementary Fig. 2a, Supplementary Fig. 3a**).

**Figure 1.**
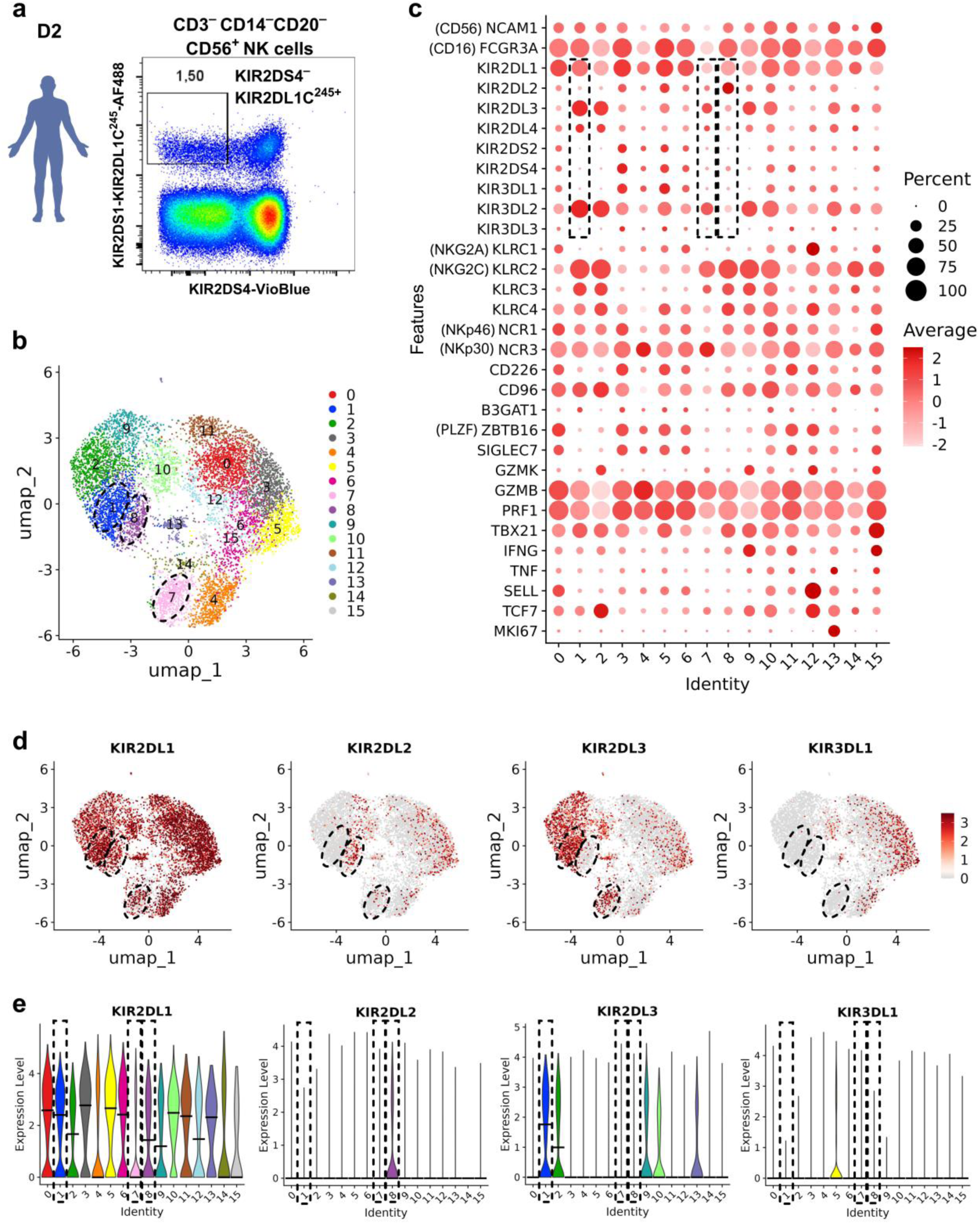
scRNA-Seq of sorted KIR2DL1C^245+^ NK cells of Donor 2. **a.** Sorting gates to obtain KIR2DL1C^245+^ NK cells. **b.** UMAP at high resolution. **c.** Dotplot with NK cell receptor genes and genes for general characterization. **d.** UMAP with selected NK cell receptor genes. **e.** Violin plots with the same genes.

We analyzed each population separately, and avoided data integration with large datasets, in order to preserve clone-specific characteristics. After quality filtering, we investigated scRNA-Seq portraits of four defined NK cell populations: 4833 KIR2DS4^+^NKG2C^+^ cells (D01), 14845 KIR2DS4^+^NKG2C^−^ cells (D01) 9104 KIR2DS4^−^KIR2DL1C^245+^ cells (D02), and 7475 KIR2DS4^+^KIR2DL1^−^ cells (D02).

For each population, we distinguished from 12 to 15 scRNA-Seq clusters (**Fig. 1b, Supplementary Fig. 1b, Supplementary Fig. 2b, Supplementary Fig. 3b**), of which 1 to 2 referred to “Resting” NK cell clusters, characterized by expression of *SELL* (CD62L, also known as L-selectin), *TCF7*, and *NKG2A*. These clusters include cells with high proliferation capacity ^48,49^, and could be associated with the NK cell *alter ego* of naive and/or central memory T cells ^50^. Each population also included “Dividing” *MKI67*^+^ cluster, and a number of clusters with distinct cytotoxicity and cytokine patterns **(Fig. 1c, Supplementary Fig. 1c, Supplementary Fig. 2c, Supplementary Fig. 3c**).

Remarkably, scRNA-Seq clusters were characterized by heterogeneous patterns of NK cell receptor expression (**Fig. 1c-e, Supplementary Fig. 1c-e, Supplementary Fig. 2c-e, Supplementary Fig. 3c-e**). This heterogeneity was particularly pronounced in scRNA-Seq clusters displaying an “adaptive” NK phenotype, defined by *KLRC1* (NKG2A)^low^, *KLRC2* (NKG2C)^high^, *ZBTB16* (PLZF)^low,^ *SIGLEC7*^low 12,17^.

In particular, on the example of KIR2DS4-KIR2DL1^+^ cells of Donor 2, scRNA-Seq Clusters 1,2,7-9, and 14 demonstrated such ‘adaptive” transcriptome profile. The distribution of KIRs among those clusters was diverse and suggests a clonal lineage. For instance, Cluster 1 showed almost no expression of *KIR2DL2*, whereas the neighboring Cluster 8 demonstrated high expression of this gene. Conversely, *KIR2DL3* showed an inverse pattern, with high expression in Cluster 1 and low/absent expression in Cluster 8 (**Fig. 1c-e**). Similar KIR receptors heterogeneity has been previously reported for clonal NK cell populations^17^.

### 2. Single NK cell culturing reveals clonally distinct receptor patterns

To further elucidate the clonal NK cell properties, we performed culturing of single-cell sorted CD56^+^CD3^−^CD20^−^CD14^−^KIR2DS4^+^ NK cells of 10 healthy donors (**Fig. 2a,b,c**). The majority of KIR2DS4^+^ NK cells belong to CD56^dim^ NK cells (**Supplementary Fig. 4a,b**), which are characterized by lower proliferation capacity compared to CD56^bright^ NK cells ^51^. The median efficiency of single NK cell culturing was ∼1% (**Fig. 2a**). 34 clones exceeding 10,000 cells after three weeks of culturing (**Fig. 2b**) were used for further flow cytometry phenotypic assessment (**Fig. 2c,d**).

**Figure 2.**
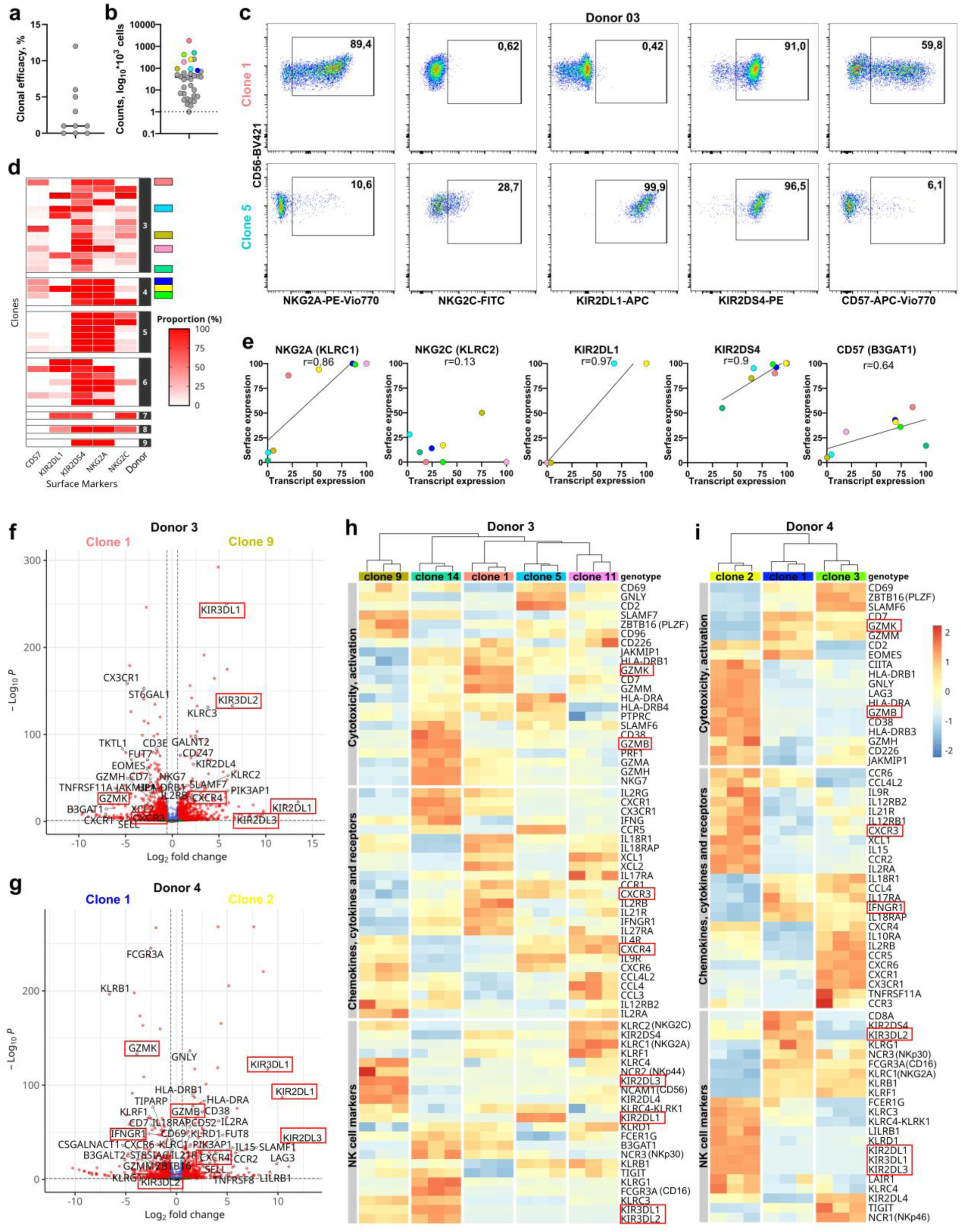
Single cell culturing of KIR2DS4^+^ NK cells. **a.** Efficiency of single cell clonal culturing, representing the percentage of clones, exceeded the 10 thousand of cells in each individual. **b.** Obtained cell counts per expanded KIR2DS4^+^ clone, presented as log10 normalized counts multiplied to one thousand. **c.** Typical flow cytometry assessment on example of Donor 3 Clones 1 and 5. **d.** Summary of major surface receptors expression assessed by flow cytometry, 34 clones of 7 donors. Proportion of marker-positive cells is shown. **e.** Correlation of normalized transcript levels with surface protein expression. Each dot represents one clone. **f,g.** Volcano plot comparing Clone 9 with Clone 1of Donor 3 and Clone 2 with Clone 1 of Donor 4, a chosen DEGs are labeled. **h,i.** Heatmap of clone-specific gene expression for three groups of NK cell genes for the five KIR2DS4^+^ clones from Donor 3 and three clones from Donor 4

In line with KIR2DS4^+^ expression on initial gated NK cell subset, all clones exhibited robust surface expression of KIR2DS4, with consistent detection across the majority of cells within each clone. About 50% of clones co-expressed the inhibitory receptor NKG2A, a feature previously linked to enhanced proliferative capacity relative to NKG2A-negative NK cells. Expression of KIR2DL1 mainly followed a binary distribution, with clones being either entirely positive or negative. In contrast, surface expression of NKG2C and CD57 varied substantially across KIR2DS4^+^ clonal cultures (**Fig. 2d**), suggesting a heterogeneous differentiation/activation/exhaustion status. To gain further molecular insights, we performed transcriptional profiling on five KIR2DS4^+^ clones from Donor D03 and three clones from Donor D04, with the highest numbers of cells. Each clone was split into 3 replicates at the level of cultured cells. Normalized transcript levels correlated with surface protein expression for most markers, except NKG2C, which showed discordant expression (**Fig. 2e**). Thousands of differentially expressed genes (DEGs) emerged from pairwise comparisons of clones’ transcriptomes (**Supplementary Fig. 1**), including multiple genes classically associated with NK cell functionality (**Fig. 2f,g**). The latter included a number of “educating” NK cell receptors associated with clonally imprinted memory^17,42^ (**Fig. 2f,g**).

In particular in Donor D03, clones specialized along distinct axes. Cytotoxic clones were defined by high effector gene expression; Clone 1 was marked by *GZMK* and *GZMM*, while Clone 14 preferentially expressed *GZMH*, *GZMB*, and *PRF1*. (**Fig. 2h**). In contrast, “helper-like” Clone 11 displayed a reverse profile: high levels of chemokines (e.g., *CCL3*, *CCL4*, *CCL5*) and cytokine receptors (*IL2RB*, *CXCR4*) coupled with low cytotoxic gene expression. Other clones showed alternative states, including an activated but low-cytokine profile (Clone 5) and a chemokine-high, low-cytotoxicity profile (Clone 9). This functional specialization was mirrored by clone-restricted patterns of education and receptor modules. Clone 9 uniquely co-expressed *KIR2DL3*, *KIR2DL4*, and *NCR2*. Clone 14 was distinguished by *KIR3DL1*, *KIR3DL2*, and *NCR3*. The highest expression of the sorting marker KIR2DS4 was observed in Clone 11, alongside increased expression of *KLRC2* (NKG2C) and *KLRC1* (NKG2A) genes. Clone 5 was defined by high *KIR2DL1* expression, consistent with phenotypic data, while Clone 1 displayed a moderate receptor profile.

A similar modular organization was observed among Donor 4 clones (**Fig. 2i**). Clone 2 preferentially expressed the cytotoxicity module, with high levels of granzymes, perforin and some activation markers, high expression of several cytokine receptor genes necessary for NK cell activation (*IL2RA*, *IL12RB1/2*, *IL21R*) and co-expressed several KIR genes, including *KIR2DL1/DL3* and *KIR3DL1*. Clone 3 showed a more balanced profile, with moderate cytotoxic gene expression but increased expression of another set of chemokine and cytokine receptors (such as *CX3CR1* and multiple interleukin receptors), and demonstrated enhanced expression of *NCR1*, *KIR2DL4* and some other genes of NK cell receptors. Clone 1, in contrast, was characterized by low expression of cytokine, chemokine and their receptor genes, and high expression of inhibitory receptor genes (e.g. *KLRG1*, *KIR3DL2*, *KLRC1*), and demonstrated the highest expression of *KIR2DS4*. Together, this analysis illustrates that cytotoxicity, migratory and receptor modules are combinatorially deployed across NK cell clones, providing a transcriptional basis for functional clonal heterogeneity.

### 3. Clonal transcriptional heterogeneity of glycosyltransferase genes

Remarkably, transcriptional diversity among KIR^+^ NK cells was not limited to classical effector and receptor genes but also extended to glycosylation pathways. In all four scRNA-Seq datasets of KIR^+^ subsets, we detected more than 30 glycosyltransferase-related genes that were differentially expressed across clusters (**Fig. 3a**). These included enzymes involved in early steps of N-glycosylation (e.g. *ALG5*, *DPAGT1*, *DPM1/2*, *TMEM258*, *OST4*, *STT3B*), branching and elongation of complex N-glycans (*MGAT4A*, *MGAT5*), O-glycan initiation (*C1GALT1*, *B3GNT7*, *B3GALNT1*), as well as fucosyl- and sialyltransferases (*FUT2*, *FUT7*, *ST3GAL2*, *ST3GAL5*). On example of KIR2DL1C^245+^ NK cells of Donor 2, previously highlighted “adaptive” clusters demonstrated the different expression of glycosyltransferase genes, where Cluster 7 demonstrated the low expression of *MGAT1* and high expression of *KRTCAP2* compared to clusters 1 and 8; Cluster 1 have shown higher expression of *MGAT4A* and lower expression of *PIGC* compared to cluster 8 (**Fig. 3b**). Similarly, KIR2DS4^+^ clusters of the same donor had a different distribution of glycosyltransferase genes (**Fig. 3c**). Each KIR^+^ subset displayed a distinct combination of these genes and, within a given subset, individual clusters showed sharply demarcated glycosyltransferase “modules”, suggesting that groups of clonally related NK cells carry stable, subset-specific glycosylation programs.

**Figure 3.**
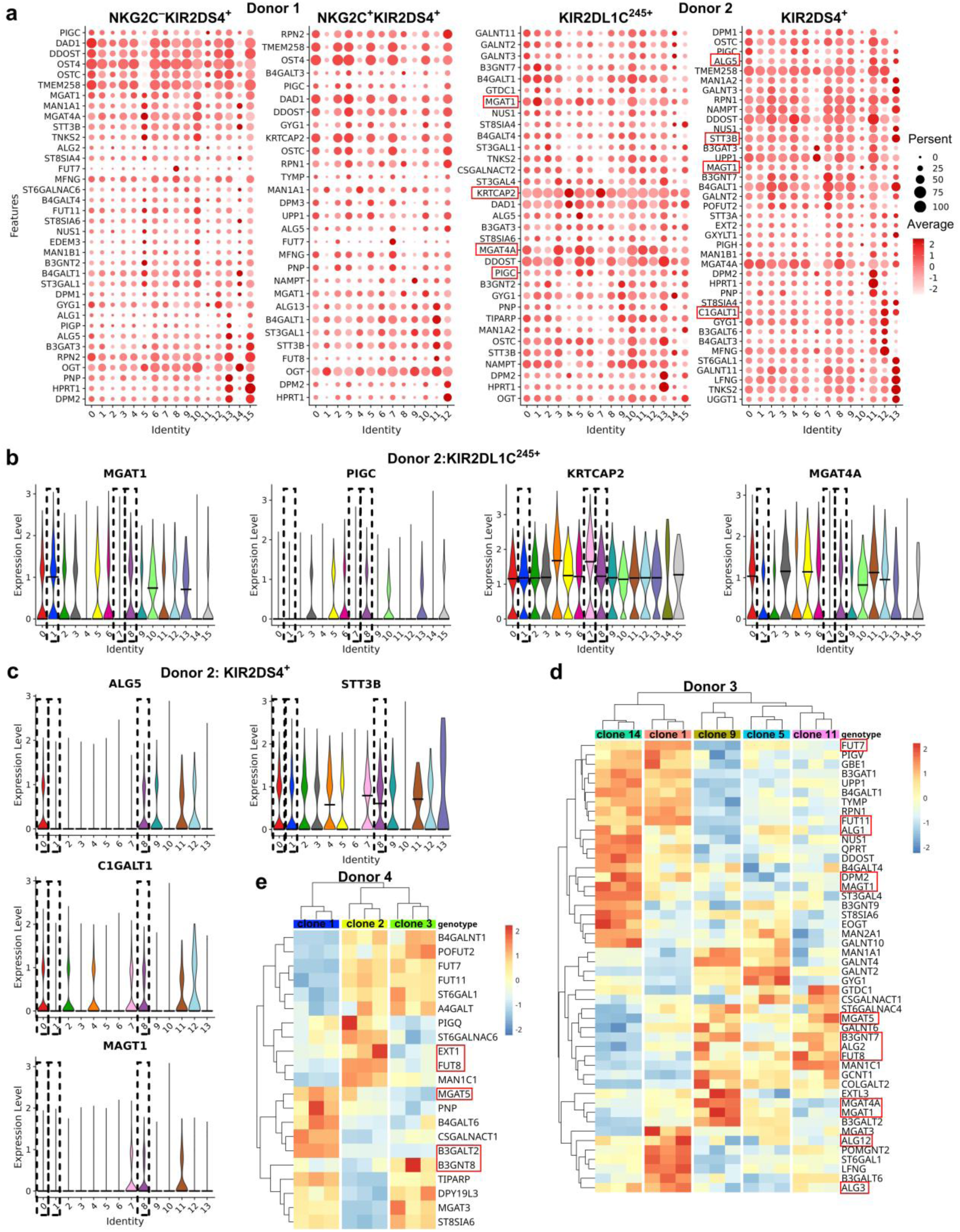
Glycosyltransferase genes expression in scRNA-Seq and Bulk RNA-Seq. **a.** Dot plots of differentially expressed glycosyltransferase genes in four scRNA-Seq datasets. **b, c.** Violin plot of selected glycosyltransferase genes for KIR2DL1+ and KIR2DS4+ NK cells of Donor 2. **d, e.** Heatmap of differentially expressed glycosyltransferase genes in Bulk RNA-Seq of for the five KIR2DS4+ clones from Donor 3 and three clones from Donor 4.

Bulk RNA-Seq of KIR2DS4⁺ single-cell-derived clonal cultures further supported the concept of clonally imprinted glycosylation programs. In Donor 3, five KIR2DS4⁺ clones showed distinct patterns of glycosyltransferase expression (**Fig. 3d**). For example, some clones were enriched in genes associated with core N-glycan synthesis (Clone 14 and Clone 1) and O-glycan synthesis (Clone 5), whereas others preferentially expressed branching MGAT genes (Clone 9), or fucosyltransferases (*FUT7* and *FUT11* - Clone 1, *FUT8* - Clone 11). A similar pattern was observed for three KIR2DS4⁺ clones from donor 4 (**Fig. 3e**), in which a different, non-overlapping subset of glycosyltransferases dominated.

Thus, both at the single-cell cluster level and in expanded clonal cultures, KIR2DS4⁺ NK cells exhibited stable, clone-specific combinations of glycosyltransferase transcripts, consistent with a clonally imprinted diversification of glycosylation machinery.

### 4. Antigen-specific culturing reveals clonal expression patterns

By analogy with antigen-specific T cell proliferation assays ^52,53^, we hypothesized that antigen-specific NK cell clones could be enriched by culturing in the presence of distinct antigenic peptides presented in a cognate MHC-I context. Presumably, during expansion, NK cell clones recognizing the target peptide may gain a proliferative advantage. Transcriptomic analysis of replicate cultures should then enable identification of reproducibly enriched transcriptional patterns characteristic of the antigen-specific clone(s).

To this end, we performed proliferation assays using CD3-depleted PBMCs to exclude potential confounding effects from memory T cells and their cytokine-mediated NK cell activation. This autologous system was previously established to identify antigen-specific NK cell responses^16,54^. CD3-depleted PBMCs were stimulated with selected peptides demonstrating binding affinity to HLA-A*11 and HLA-C*06 (**Table 2**).

**Table 1.** Donors’ characteristics.

| No | Gender | Age | hCMV<br>IgG | EBV<br>IgG | SARS-CoV-2 IgG | HLA-A | HLA-B | HLA-C |
| --- | --- | --- | --- | --- | --- | --- | --- | --- |
| D1 | Female | 33 | 5.1 | 293 | 3,75 | A*02:01:01<br>A*11:01:01 | B*35:01:01<br>B*57:01:01 | C*04:01:01<br>C*07:01:01 |
| D2 | Male | 29 | 2.1 | 0 | 27,6 | A*29:02:01 | B*44:03:01<br>B*45:01:01 | C*06:02:01<br>C*16:01:01 |
| D3 | Male | 25 | 0.3 | 180 | 52,7 | A*02:01:01<br>A*03:01:01 | B*35:01:01<br>B*51:01:01 | C*04:01:01<br>C*15:13 |
| D4 | Male | 45 | 1.3 | 130 | 6,4 | A*02:01:01<br>A*02:05:01 | B*18:01:01<br>B*50:01:01 | C*06:02:01<br>C*07:01:01 |
| D5 | Female | 41 | 8.3 | 0 | 33,7 | A*26:01:01<br>A*29:02:01 | B*44:03:01<br>B*49:01:01 | C*07:01:01<br>C*16:01:01 |
| D6 | Female | 21 | 2.4 | 90 | 7,7 | A*02:01:01<br>A*03:01:01 | B*07:02:01<br>B*08:01:01 | C*07:01:01<br>C*07:02:01 |
| D7 | Female | 61 | 8 | 268 | 27,5 | A*26:01:01<br>A*30:01:01 | B*35:02:01<br>B*55:01:01 | C*03:03:01<br>C*06:02:01 |
| D8 | Male | 22 | 0.8 | 198 | 16,1 | A*02:01:01<br>A*11:01:01 | B*35:03:01<br>B*40:02:01 | C*02:02:02<br>C*04:01:01 |
| D9 | Male | 71 | 0 | 0 | 35,7 | A*24:02:01 | B*07:02:01<br>B*41:01:01 | C*07:02:01<br>C*16:04:01 |
| D10 | Female | 26 | 0 | 0 | 9,7 | A*11:01:01<br>A*26:01:01 | B*07:02:01 | C*07:02:01 |

**Table 2.**
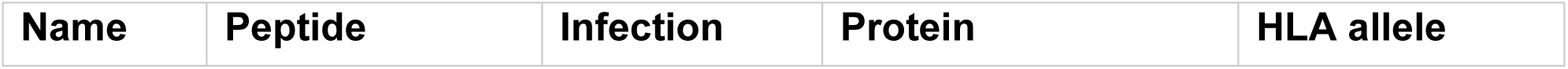
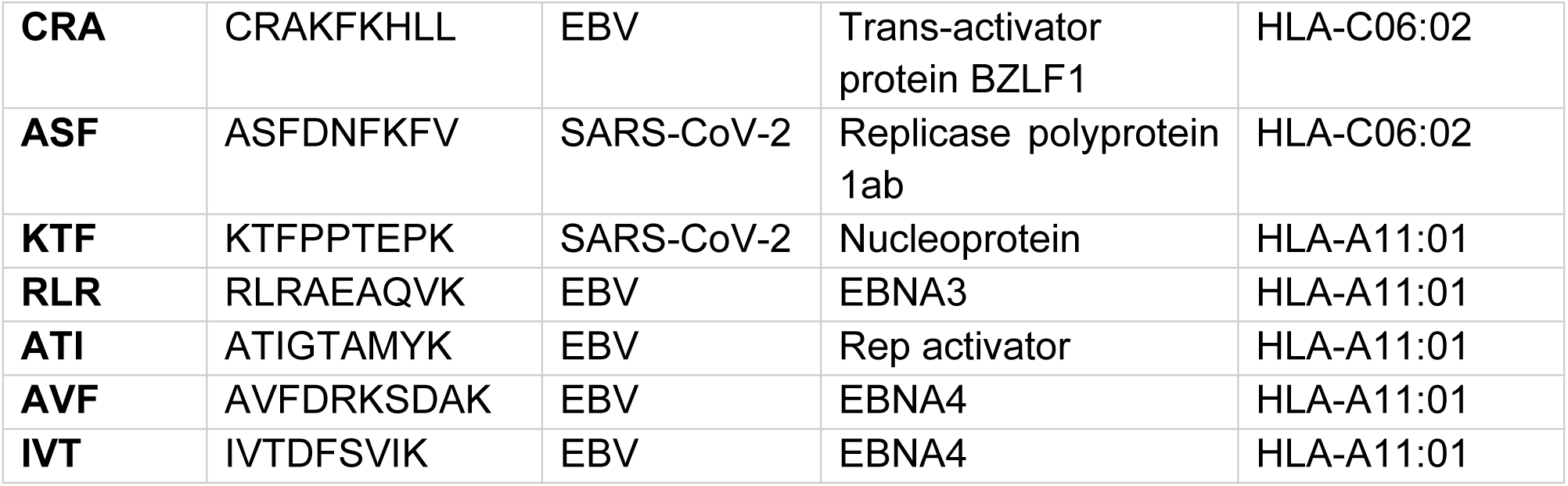
List of peptides.

We performed two antigen-specific proliferation experiments on PBMC of HLA-A*11-positive Donor 1, whose NK cells were profiled by scRNA-Seq (see above). In order to identify transcriptional imprints of reproducibly expanding NK cell clones, proliferation assay for each antigenic stimulus was performed in three independent replicates. Proliferated CFSE^low^ NK cells were FACS-sorted for transcriptomic analysis (**Fig. 4a**).

**Figure 4.**
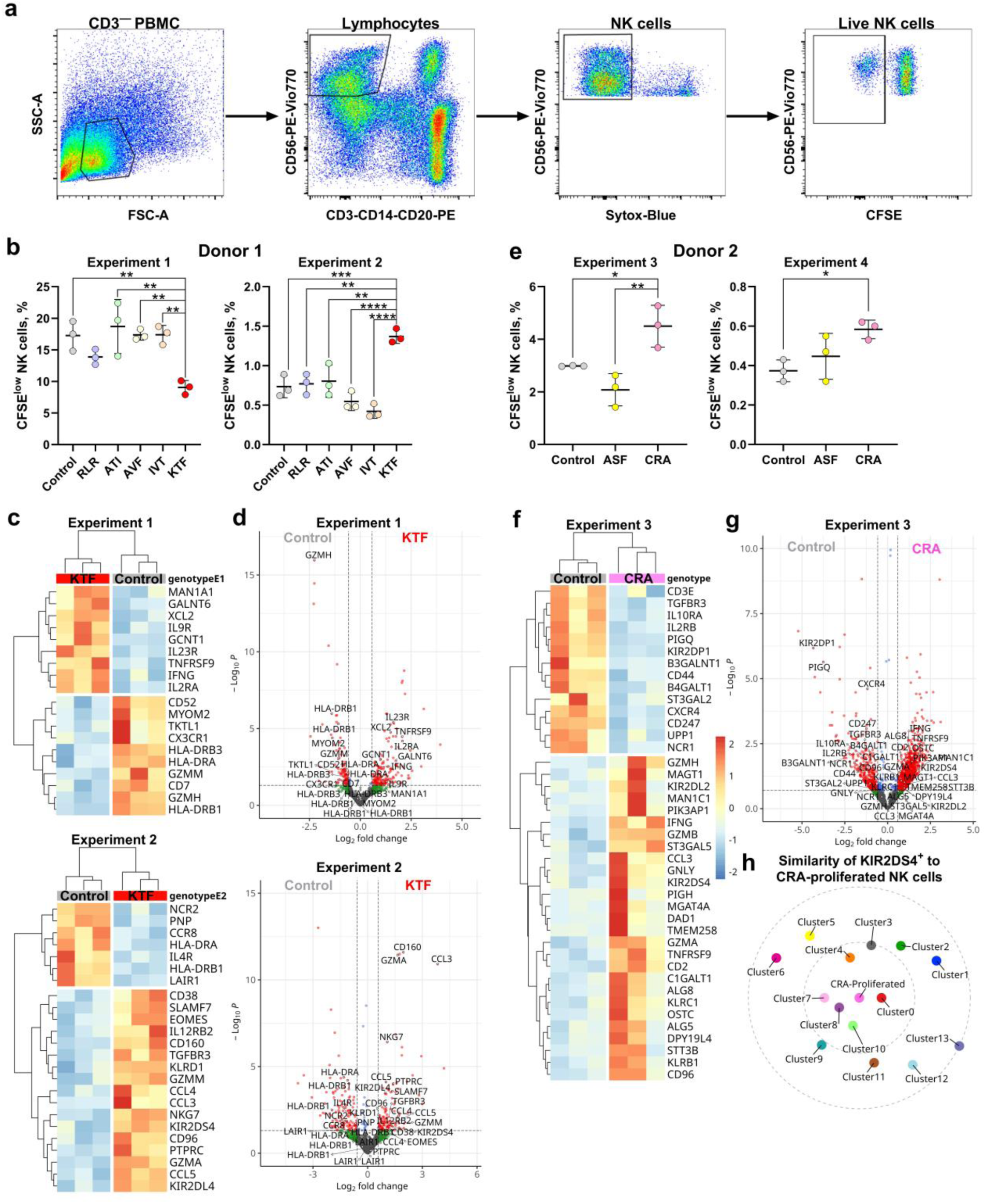
Antigen-specific proliferation of NK cells. **a.** Gating scheme of obtaining CD56^+^CD3^−^CD20^−^CD14^−^SYTOX^−^CFSE^low^ NK cells. **b.** Proportion of CFSE^low^ NK cells stimulated with peptides loaded in HLA-A11 of Donor 1 in two Experiments. **(c.** Heatmaps of selected DEGs between KTF-proliferated and control-proliferated NK cells of Donor 1 in two Experiments. **d.** Volcano plot comparing KTF-proliferated and control-proliferated NK cells of Donor 1 in two Experiments. **e.** Proportion of CFSE^low^ NK cells stimulated with peptides loaded in HLA-AC06 of Donor 2 in two Experiments. **f.** Heatmap of selected DEGs between CRA-proliferated and control-proliferated NK cells of Donor 2 in Experiment 3. **g.** Volcano plot comparing CRA-proliferated and control-proliferated NK cells of Donor 2 in Experiment 3. **h.** Distance plot of KIR2DS4^+^ scRNA-Seq to CRA-proliferated NK cells.

Unexpectedly, in the first proliferation assay (Experiment 1), stimulation with the SARS-CoV-2 peptide KTFPPTEPK (KTF) specifically resulted in a marked inhibition of NK cell proliferation as validated by CFSE assay compared to control and other peptide-stimulated samples^55^ (**Fig. 4b**). On the contrary, a repeat assay performed three months later for the same donor (Experiment 2) yielded the opposite results, with KTF stimulation inducing significant NK cell proliferation relative to peptide-free controls and samples stimulated with other peptides (**Fig. 4b**). Notably, baseline proliferation in control samples declined substantially in Experiment 2 compared to Experiment 1. As other peptides elicited no proliferation differences in either experiment, we focused transcriptomic analyses on proliferated NK cells from control and KTF-stimulated conditions across both timepoints.

In Experiment 1, NK cells proliferating in the presence of KTF peptide were characterized by a higher expression of transcripts encoding key cytokine gene *IFNG*, cytokine receptor genes *IL2RA*, *TNFRSF9* (4-1BB or CD137), *IL23R*, *IL9R*, and glycosyltransferase genes *MAN1A1*, *GALNT6*, and *GCNT* compared to the control cultures. Conversely, genes encoding granzymes *GZMM* and *GZMH*, MHCII genes *HLADRA* and *HLADRB1/3*, and the intercellular protein gene *MYOM2* (markers enriched in classical NK cells ^56^, were expressed at lower levels (**Fig. 4c,d**).

In Experiment 2, NK cells proliferating in the presence of KTF were characterized by higher expression of chemokine genes *CCL3*, *CCL4*, *CCL5*, activation and cytotoxicity-associated genes *CD38*, *PTPRC*, *GZMM*, *GZMA*, *NKG7*, MHCI-interacting receptor genes *CD160*, *KIR2DL4* and *KIR2DS4* (**Fig. 4d**). Low expressed genes included NCR2, cytokine and chemokine receptors *IL4R* and *CCR8*, and class MHCII genes (*HLADRA*, *HLADRB1*).

The possible explanation for observed differences is that in the two distinct time points, distinct NK cell clones dominated in antigen-specific response to the KTF peptide. In Experiment 1, responding clone(s) could have suppressive characteristics while clone(s) proliferated in Experiment 2 demonstrated a cytotoxically active profile. Regardless of different proliferation outcomes in these experiments, the selective action of the peptide, presented in the appropriate context, supports a distinct peptide-specific clonal-organized response.

Addressing the same issue, the NK cell proliferation assay was applied for Donor D02, who has the HLA-C*06 genotype. The CRAKFKHLL (CRA) EBV-derived peptide and ASFDNFKFV (ASF) SARS-CoV-2-derived peptide were selected for testing. Similar to Donor 1 experiments, Experiments 3 and 4, conducted with a 3-month gap between them, showed different levels of baseline NK cell proliferation. However, we observed enhanced NK cell proliferation in response to the CRA peptide in both experiments (**Fig. 4f**).

Experiment 3 revealed over a thousand differentially expressed genes, indicating a possible clonal NK cell response specific to EBV. Similar to Experiment 2, NK cells proliferated in the presence of the CRA peptide demonstrated enhanced expression of receptor genes *KIR2DL2* and *KIR2DS4* along with several granzyme genes. Besides, increased transcript levels of *GNLY*, *CD2*, and glycosyltransferase genes *MAGT1*, *MAN1A1*, *ST3GAL5*, *STT3B*, *OSTC*, *C1GALT1*, *ALG5/8*, and *MGAT4A* were observed. Genes with decreased expression included genes of cytokine and chemokine receptors *CX3CR1*, *TGFBR3*, *IL2RB*, *IL10RA*, and *CXCR4*; the *NCR1* receptor gene and glycosyltransferase genes *ST3GAL2*, *B3GALNT1*, *FUT2*, *PIGQ* and *B4GALT1* (**Fig. 4g,h**). In Experiment 4, the low counts of proliferating NK cells did not allow reliable RNA-Seq analysis.

We next asked whether glycosyltransferases that were identified during antigen-driven proliferation corresponded to any pre-existing transcriptional modules in KIR2DS4^+^ NK cells (**Fig. 4i**). Donor D02 NK cells cultured in presence of the EBV peptide CRA were characterized by enlarged expression of *KIR2DS4*, *KIR2DL2*, *MAGT1*, *MAN1A1*, *ST3GAL5*, *STT3B*, *OSTC*, *C1GALT1*, *ALG5/8* and *MGAT4A* genes, with concomitant decrease in expression of *ST3GAL2*, *B3GALNT1*, *FUT2*, *PIGQ* and *B4GALT1*. In the KIR2DS4^+^ scRNA-Seq dataset from the same donor, three clusters demonstrated the co-expression of KIR2DL2 with KIR2DS4 (clusters 0, 4, and 8 on **Supplementary Fig. 1**). Several of these genes (*ALG5*, *C1GALT1*, *MAGT1*, *STT3B*, *MGAT4A*, *TMEM258*) were not uniformly expressed and demonstrated the most similarity within cluster 8 (**Fig. 3d**), indicating that CRA peptide could potentially expand a pre-existing subset of clonal KIR2DS4⁺ NK cells with a characteristic glycosyltransferase signature.

In summary, these experiments demonstrate that antigen-specific proliferation enables the enrichment of distinct NK cell clones responsive to particular peptides. The proliferating populations are distinguished by specific profiles of characteristic NK cell receptors, lymphoid receptors, cytokines, and glycosyltransferase genes. We propose that these patterns correspond to distinct clonal NK cell populations bearing defined effector programs and patterns of activation/inhibition of attack, including patterns of specific or semi-specific antigen recognition.

### 5 Post-translational glycosylation modulates KIR2DS4-CRA-HLA-C*06 interactions *in silico*

Given the observed diversity of glycosyltransferase expression among antigen-specific NK cell clones, we hypothesized that post-translational modifications, particularly glycosylation, may influence the structure and ligand-binding dynamics of KIR receptors. KIRs are known to undergo both phosphorylation and glycosylation. To investigate the location of N-linked and O-linked glycosylation sites, we used the bioinformatic service^57^, which predicts the possibility of each glycosylation to occur. The complexes of KIR2DL1, KIR2DL2, KIR2DL3, KIR2DS2 and KIR3DL1 with corresponding HLA ligands were available in PDB crystal structures and were analyzed (**Fig. 5a**). Each KIR has shown several N-linked glycosylation sites with high binding potential (higher than 0.5) and at least one O-linked glycosylation site (**Fig. 5a**). One N-glycosylation site of KIR2DL2/DL3/DS1 was located exactly above the peptide; the probable glycosylation on this site could influence the KIR-peptide interaction. At least one more N-linked site in all KIR located in close proximity within the binding interface.

**Figure 5.**
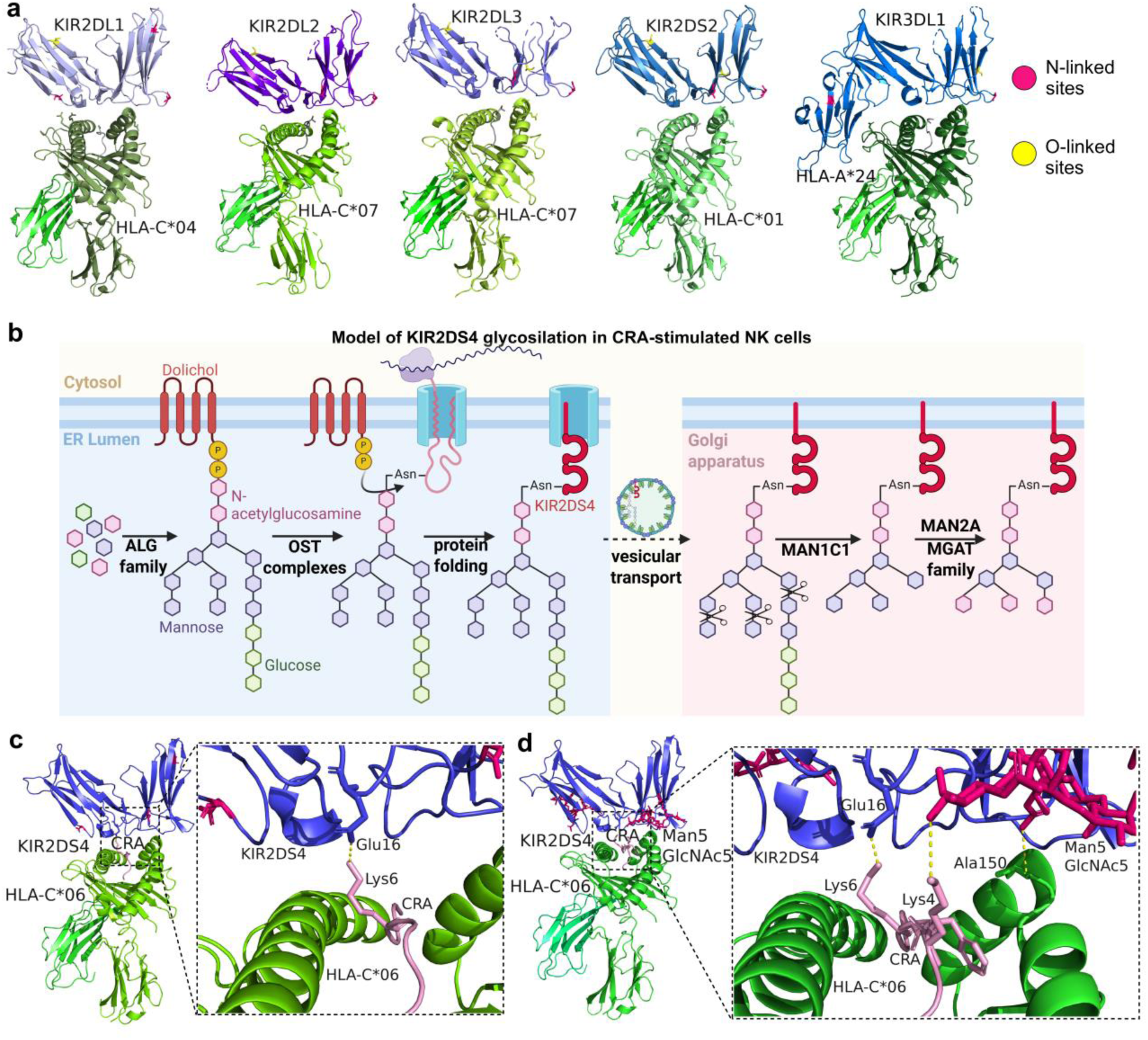
Structures of KIR-HLA complexes with peptides. **a.** Representation of N- and O-linked glycosylation sites for different crystal structures of KIR-peptide-HLA complexes. **b.** Model of KIR2DS4 glycosylation based on increased expression of glycosyltransferase genes in CRA-proliferated NK cells. **c,d.** The AlphaFold 3 prediction of KIR2DS4–CRA–HLA-C*06 complexes depending on the glycosylation type: without modification (c) and with Man_5_GlcNAc_5_ modification at Asn41 and Asn185 (d).

Recent work has demonstrated that N-linked glycosylation of domain 0 in KIR3DL1 modulates allele-specific HLA interactions, suggesting that glycosylation can fine-tune receptor–ligand binding affinity and specificity^40^. In CRA-stimulated NK cells of Donor D02 (as well as in scRNA-Seq cluster 8) we identified the increased expression of genes which encodes the proteins of OST-A and OST-B complexes (*STT3B*, *MAGT1*, *OSTC*, *DAD1*, *TMEM258*)^58^. This complex can catalyze the N-linked glycosylation of transmembrane proteins^59^ mostly by the Glc_3_Man_9_GlcNAc_2_ addition^60^. However, in some cases OST complexes can transfer incomplete glycans^61^. The preassembly of the glycans is mainly conducted by ALG glycosyltransferase protein group^60^, from which the *ALG5* and *ALG8* gene expression was enhanced increased in NK cells proliferated in presence of CRA peptide. After the addition of initial glycan in ER, the glycosylated proteins could be remodeled in the Golgi Apparatus. Mannosidase Alpha Class 1C Member 1 encoded by *MAN1C1* may conduct trimming of a specific mannose sugars, converting the high-mannose structure into a smaller form, such as Man_5_GlcNAc_2_^62^. Further, high expression of *MGAT4A*, may form a β1-4 GlcNAc branch on the α1-3 mannose arm in N-glycans^63^ (**Fig. 5b**). In addition to N-linked glycosylation the increased expression of *C1GALT1* may act on any serine or threonine residues that have been previously modified with a single GalNAc sugar and convert these simple modifications into the O-glycan structure (Galβ1-3GalNAc-)^64^.

Even though we could not provide the evidence that those enzymes are exactly glycosylase the KIRs, we can explore this possibility for KIR2DS4. We first identified the predicted N- and O-linked glycosylation sites on the receptor^57^, and subsequently modeled the KIR2DS4–CRA–HLA-C*06 complex using AlphaFold3^65^. We previously validated the structural accuracy of AlphaFold predictions for KIR2DS4^66^. Five N-linked glycosylation sites were identified, from which three demonstrated the binding potential higher than 0.5, all possible O-linked sites had the potential lower than 0.5, where the highest potential equal to 0.48 was reached for Thr112. Two predicted N-linked glycosylation sites at Asn41 and Asn185 were located in close proximity to the HLA-binding interface and were further modified.

In the unmodified complex, the KIR2DS4 receptor formed one polar contact with the CRA peptide: Lys6 of the peptide engaged Glu16 on KIR2DS4 (**Fig. 5c**). To evaluate the structural consequences of glycosylation, we modified Asn41 and Asn185 with a branched Man_5_GlcNAc_5_. which could be potentially formed based on the high expression of glycosyltransferase genes in CRA-proliferated NK cells (**Fig. 5b**). In addition to the contact of Lys6 of the peptide and Glu16 on KIR2DS4 the N-acetyl-β-D-glucosamine of Asn185 modification has formed the contact with Lys4 of the CRA peptide (**Fig. 5d**). Moreover, the Man_5_GlcNAc_5_ as well demonstrated the formation of contact with Ala150 of HLA-C*06 (**Fig. 5d**).

Collectively, these structural simulations suggest that site-specific glycosylation of KIR2DS4 can substantially modulate both receptor-ligand and receptor-peptide interactions, potentially influencing antigen recognition by NK cells.

## Discussion

The clonal organization of NK cells has emerged as a fundamental feature of their biology, with profound implications for immune memory and functional diversification^8,9,17,24,67^. Our data extend the concept of adaptive and clonally expanded NK cells, which has been mostly defined in the context of CMV-driven NKG2C⁺ expansions, to KIR⁺ NK-cell clones bearing diverse receptor, effector and migratory programs. Strikingly, KIR-expressing NK cells exhibit not only the expected heterogeneity in cytotoxicity and receptor modules, but also sharply demarcated, stable glycosyltransferase expression programs that differ between clones. This constitutes a previously unrecognized post-translational layer of NK cell diversification. These observations raise several important considerations for understanding NK cell clonality and its functional consequences.

The best-characterized model of adaptive NK cell memory centers on NKG2C^⁺^ CD57^⁺^ NK cells that expand preferentially in response to hCMV infection and acquire a stable epigenetic and transcriptional imprint^12,15,17,68^. Lineage-tracing using mitochondrial DNA mutations has confirmed that many NK cell subsets in adult humans represent long-lived clonal lineages^41^, and retroviral barcoding in both murine^8^ and non-human primate models^9^ has established that antigen-driven clonal expansion, rather than polyclonal proliferation, is the dominant mode of NK cell memory formation. Our scRNA-Seq analysis of KIR2DS4^⁺^ and KIR2DL1^⁺^ NK cell populations revealed transcriptionally distinct clusters per sorted subset, each displaying a characteristic combination of KIR, KLRC, and NCR receptor genes. The inverse distribution of KIR2DL2 and KIR2DL3 between neighboring clusters is consistent with the stochastically chosen but subsequently clonally stable KIR expression model established in previous works^43,69^. Importantly, clusters displaying the “adaptive” transcriptional phenotype (*KLRC1*^low^, *KLRC2*^high^, *ZBTB16*^low^, *SIGLEC7*^low^)^17,68^ showed the most pronounced KIR heterogeneity, consistent with the expectation that differentiated, antigen-experienced clones occupy distinct receptor niches.

Bulk RNA-seq of single-cell-derived KIR2DS4^⁺^ clonal cultures provided independent and complementary evidence for functional clonal specialization. Clones from the same donor were segregated into recognizable functional modules: cytotoxic clones enriched in *GZMB*, *GZMH*, and *PRF1*; chemokine-secreting clones expressing *CCL3*, *CCL4*, and *CCL5*; and receptor-enriched clones co-expressing distinct combinations of inhibitory and activating KIRs. This modular organization mirrors the functional diversity described for NKG2C^⁺^ adaptive NK cell clones^17^ and extends it to KIR2DS4^⁺^ populations, demonstrating that clonal functional specialization is not an exclusive feature of the NKG2C-mediated adaptive NK cell axis.

The diverse granzyme co-expression profiles across KIR2DS4^⁺^ clones are of particular interest. We identified typical patterns of distribution of these effector molecules in proliferating NK cells and characterizes both types of clonal cultures as combining several different mechanisms of cell death induction. Clone 1 from Donor D03 was characterized by preferential expression of *GZMK* and *GZMM*, whereas Clone 14 expressed predominantly *GZMB* and *GZMH*. Granzyme K, through its trypsin-like activity (by cleaving substrates after Arg or Lys), induces mitochondrial ROS production and chromatin condensation. This is combined with the action of Granzyme M, which exhibits atypical specificity (cuts after Met or Leu) and induces death independently of caspases^70^. The clone 14 expresses the widespread cytolytic Granzyme B along with Granzyme H, which has chymotrypsin-like activity (Phe or Tyr) and is capable of inactivating proteins involved in adenovirus assembly and replication and/or inhibiting Granzyme B and cleaving cellular phosphoprotein La, which is necessary for hepatitis C virus (HCV) replication^71^. Thus, we have demonstrated the diversity of KIR2DS4^+^ NK cell clones, some of which may be involved to a greater extent in the proinflammatory immune response, while others are involved in cytotoxic antiviral reactions analogous to memory CD8^⁺^ T cell differentiation.

Among the molecular markers of adaptive NK cell differentiation, NKG2C (encoded by KLRC2) occupies a central position, as NKG2C^⁺^CD57^⁺^ NK cells are considered the canonical adaptive NK cell population in hCMV-seropositive individuals^12,13,15^. In our clonal culture system, we observed a systematic discordance between *KLRC2* transcript levels and NKG2C surface protein expression suggesting complex mechanisms for the regulation of NKG2C expression on the cell surface. The primary mechanism for stabilizing NKG2C at the cell surface is its assembly with the signaling adaptor DAP12, whose expression is necessary for transport of the NKG2C/DAP12 complex to the plasma membrane^72^. We did not observe consistent differences in *TYROBP* (DAP12) transcript levels between clones with discordant NKG2C mRNA and protein expression, suggesting that DAP12 availability is not the only limiting factor in these cultures. Preliminary observations from our group indicate that NKG2C surface expression may be dynamically acquired or lost during prolonged clonal culture, with loss predominating in NKG2C^⁺^ clones derived from CD56^dim^ CD57^−^ progenitors but not from CD57^+^ highly differentiated NK cells, suggesting that terminal differentiation imparts additional stability to NKG2C surface expression^73^. These observations challenge the interpretation of NKG2C surface staining as a static, lineage-defining marker.

A central and unexpected observation is that KIR⁺ NK-cell clones exhibit sharply demarcated, stable glycosyltransferase modules, both at the single-cell cluster level and in expanded KIR2DS4⁺ clonal cultures. Glycosylation is known to shape NK receptor function and ligand recognition; for instance, through modulation of KIR3DL1-HLA, 2B4-CD48, CD16-IgG, NKp30-B7-H6 interactions^36–38,40^. Our data suggest that entire glycosylation pathways, including early N-glycan assembly (ALG family, OST complex components), branching (MGAT4/5), fucosylation, and O-glycan initiation, can be clonally imprinted and may partially maintained over time, effectively creating clone-specific “glycosylation repertoires” in human NK cells. The consistency of glycosyltransferase modules both in scRNA-Seq clusters and in expanded bulk cultures strongly argues against these patterns reflecting random transcriptional noise and instead supports stable, clonally maintained glycosylation programs. While the epigenetic mechanisms underlying such maintenance remain to be formally demonstrated, precedent exists for stable epigenetic encoding of receptor expression programs in adaptive NK cell clones^17^.

Notably, different clones showed enrichment of distinct glycosylation pathways: some were characterized by elevated expression of genes involved in the core N-glycan assembly complex (OST-A/B components STT3B, MAGT1, OSTC, DAD1, TMEM258), while others preferentially expressed branching enzymes (MGAT4A, MGAT5) or fucosyltransferases (FUT7, FUT11, FUT8). This pathway-level heterogeneity is significant because it implies not only quantitative differences in glycosylation between clones, but qualitative differences in glycan structure.

Antigen-driven proliferation assays demonstrated that defined viral peptides presented by autologous MHC class I molecules can selectively enrich distinct NK cell clonal populations. The EBV-derived peptide CRAKFKHLL (CRA), presented in an HLA-C*06:02 context, reproducibly induced NK cell proliferation in Donor 2 across two independent experiments separated by three months. The transcriptional signature of CRA-expanded cells was consistent across experiments and included elevated expression of *KIR2DL2*, *KIR2DS4*, cytotoxicity genes (*GZMA*, *GZMB*, *GZMH*, *GNLY*), and a specific subset of glycosyltransferase genes (*ALG5, C1GALT1, MAGT1, STT3B, MGAT4A*). The correspondence between this transcriptional profile and that of cluster 8 in the KIR2DS4^⁺^ scRNA-Seq dataset from the same donor suggests that antigenic stimulation selectively expands a pre-committed clonal subset, defined by both a characteristic KIR repertoire and a distinct glycosylation machinery, rather than triggering a de novo transcriptional reprogramming. This is conceptually analogous to the selective expansion of pre-existing antigen-specific T cell clones following rechallenge^53^, and supports the view that antigen-specific NK cells maintain a stable, clonally encoded transcriptional identity across their lifespan.

In contrast to the consistent CRA response, the SARS-CoV-2 nucleoprotein peptide KTFPPTEPK (KTF), presented in an HLA-A*11:01 context, elicited markedly different, and in one instance opposite, responses in the same donor at two different timepoints. In Experiment 1, KTF stimulation resulted in relative suppression of NK cell proliferation accompanied by transcriptional features consistent with an immunosuppressive or regulatory NK cell phenotype. Three months later, in Experiment 2, KTF stimulation induced robust proliferation of cells with a cytotoxic, chemokine-secreting phenotype. The possible explanation for these opposing outcomes is that distinct NK cell clones dominated the KTF-specific response at the two timepoints, with different effector programs encoded within each clone. During the three-month period the donor’s inflammatory and infectious milieu may have shifted, which could have altered the clonal composition of the peripheral NK cell pool, as has been described in the context of hCMV reactivation and other inflammatory triggers^35,74,75^. Such dynamic reshaping of the antigen-specific NK cell repertoire has been proposed by analogy with T cell immunodominance hierarchies, where the relative representation of epitope-specific clones fluctuates with antigenic history and immune competition^76^. This temporal variability underscores the importance of longitudinal sampling and replicated proliferation assays in studies of NK cell antigen-specific immunity, and suggests that the functional heterogeneity of the NK cell response to a given peptide reflects the clonal composition of the KIR repertoire at the time of stimulation rather than an intrinsic property of the peptide–receptor interaction.

The observation that both EBV and SARS-CoV-2 peptides induced clonal NK cell responses is consistent with our previous demonstrations of KIR2DS4 involvement in NK cell responses to these pathogens^54,77^. The differential roles of HLA-A*11:01 and HLA-C*06:02 as restriction elements for KIR2DS4 recognition are also notable: while HLA-C*05:01-restricted KIR2DS4 binding has been the best-characterized interaction^34^, and HLA-A*11:01 interaction has been proposed based on structural homology with KIR3DL2^44^, our functional data provide in vitro evidence that peptides presented by both HLA-A*11:01 and HLA-C*06:02 can drive selective proliferation of KIR2DS4^⁺^ NK cell clones, broadening the recognized HLA restriction of this activating receptor.

Structural modeling of the KIR2DS4-CRA-HLA-C*06:02 complex identified potential N-linked glycosylation sites on KIR2DS4, located in close proximity to the HLA-binding interface. Compared to unmodified model, modification of Asn185 with a Man5GlcNAc2 core glycan introduced an additional contact between the terminal GlcNAc residue and Lys4 of the CRA peptide, and also formed a contact with HLA-C*06:02, expanding the receptor-peptide-HLA contact network. The transcriptional and structural data support a coherent glycan biosynthetic pathway operating in antigen-expanded KIR2DS4^⁺^ NK cell clones that could plausibly produce the glycan structures modeled at the KIR2DS4 binding interface. Even though these structural conclusions are based on computational modeling and not on direct biochemical characterization of KIR2DS4 glycoforms from primary NK cells; the concordance between the transcriptionally inferred glycosylation program of antigen-expanded clones and the structural modeling results establishes a strong hypothesis that site-specific glycosylation of KIR2DS4 may constitute a previously unrecognized determinant of peptide-selective recognition.

In summary, the present study establishes that KIR^⁺^ human NK cells comprise clonally imprinted populations defined by characteristic combinations of receptor, effector, and glycosyltransferase expression programs. Single-cell and clonal transcriptomics converge on the conclusion that glycosylation pathway diversity is a stable feature of NK cell clonal identity, representing a previously unrecognized post-translational dimension of the NK cell clonal repertoire. Antigen-specific proliferation assays demonstrate that viral peptides can selectively enrich pre-committed clonal subsets bearing defined receptor and glycosylation profiles, and that the identity of the responding clone may vary dynamically with the immunological history of the individual. Structural modeling provides an initial mechanistic framework in which site-specific glycosylation of KIR2DS4 expands the receptor-peptide contact network, suggesting that clonally imprinted glycosylation programs may contribute to the peptide selectivity of activating KIR-mediated recognition. These findings advance our understanding of NK cell clonal biology and suggest glycosyltransferase diversification as a novel mechanism through which the NK cell repertoire could achieve functional heterogeneity and antigen responsiveness.

## Materials and methods

### Samples

Peripheral blood samples were obtained from healthy volunteers. The study was conducted in accordance with the Declaration of Helsinki and approved by the local Ethics Committee of Pirogov Russian National Research Medical University (protocol #252 from 25 June 2025).

To isolate the peripheral blood mononuclear cells (PBMC), blood samples were collected in EDTA-containing test tubes and centrifuged in a Ficoll gradient (PanEco, Moscow, Russia). Serum samples were collected in tubes with gel-clot activators and stored at −60 °C until used. PBMCs further underwent the positive magnetic separation of CD3^+^ cells (RWD Life Science, China) to obtain CD3-depleted PBMC with a purity of less than 1% T cells in lymphocytes, or negative magnetic separation of NK cells (RWD Life Science, China) to obtain pure NK cells (98%). The resulting CD3-depleted PBMCs were cultured in the following medium: DMEM (PanEco, Russia) and NK-MACS Medium (Miltenyi Biotec, Bergisch Gladbach, Germany) in a 1:1 ratio, supplemented with 2 mM L-glutamine, 1 mM sodium pyruvate, 10% fetal bovine serum (FBS, HyClone, USA), 1% antibiotic (Antibiotic-Antimycotic Solution, Corning, USA) hereinafter referred to as complete medium. Cells were cultured in a 96-well round-bottom plate in a CO_2_ incubator at 37°C. Pure NK cells were immediately sorted to perform scRNA-Seq analysis or clonal culturing.

### Peptides

For the selection of SARS-CoV-2 and EBV peptides, peptides that bind with high affinity to HLA-C*06 or HLA-A*11 were selected based on available repositories. The list of peptides is presented in **Table 2**. Peptides were synthesized by GenScript (USA).

### Detection of hCMV- and EBV-Specific Antibodies

EBV anti-EBNA-1 IgG, SARS-CoV-2 IgG, and hCMV-IgG titers were determined via enzyme-linked immunosorbent assay using a commercial kit in accordance with manufacture instructions (Vector-Best, Novosibirsk, Russia).

### Flow cytometry

The analysis of surface expression markers with fluorescently labeled monoclonal antibodies was performed using a Longсyte C3080 equipped with 405 nm, 488 nm, and 635 nm lasers (Challenbio, China) or FACSVantage DiVa cell sorter equipped with lasers: 405, 488, 590, and 633 nm (BD BioSciences, USA). The detailed information about antibodies is presented in Table 3.

**Table 3.**
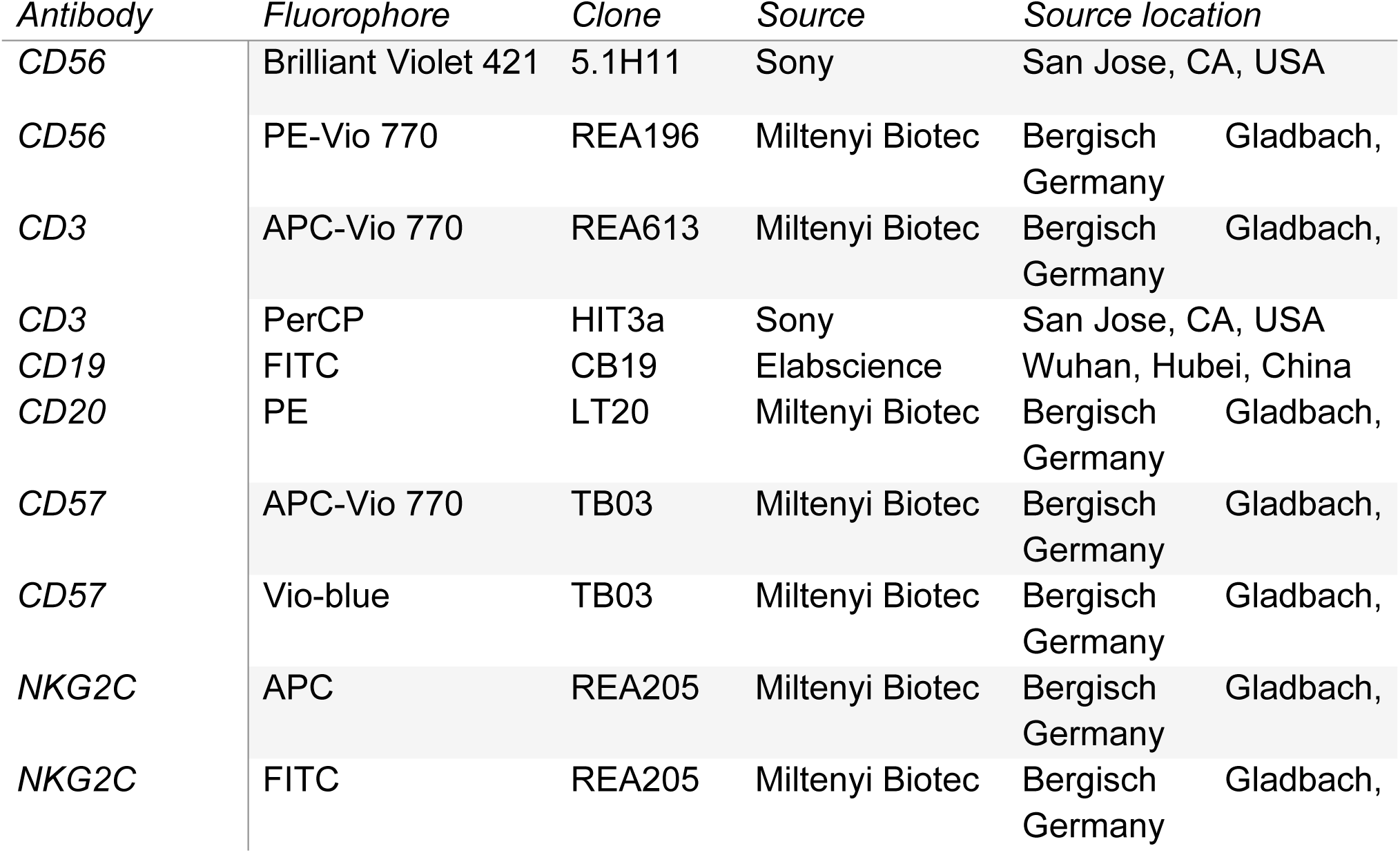
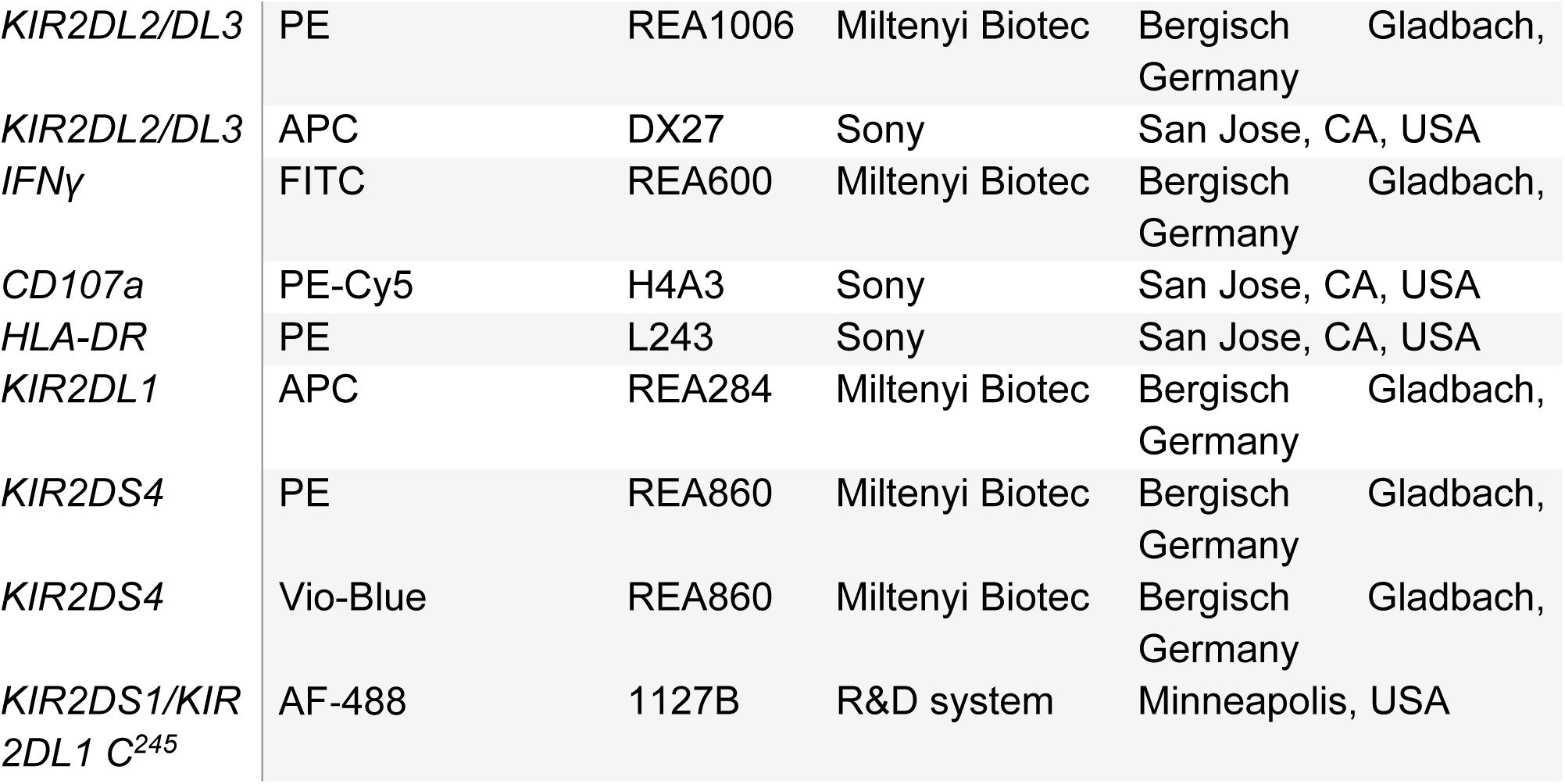
Anti-human antibody list.

### Preparation of libraries for HLA-genotyping

PBMC samples were lysed using ExtractRNA (Evrogen, Moscow, Russia). Total RNA was extracted using standard chloroform/isopropanol isolation instruction. First-strand cDNA synthesis was performed using sequence-specific primers targeting highly conserved regions of the HLA-A, HLA-B, HLA-C, HLA-DQB1, and HLA-DRB1 genes. Amplification of the cDNA was then carried out in eight separate PCR reactions per donor, employing primer mixes designed to cover exons 2–4 of HLA class I genes (HLA-A, -B, -C) and exons 2–3 of HLA class II genes (HLA-DQB1, -DRB1). Eight products of the first PCR reactions for each donor were mixed and subjected to a second amplification round using the IDT for Illumina UD Indexes kit (Illumina, USA). Library purification was performed with AMPure XP beads (Beckman Coulter, USA) following the manufacturer’s protocol. Final libraries were sequenced using 250 bp paired-end sequencing on Illumina MiSeq platform (Illumina, USA).

### Analysis of HLA-genotype

Bioinformatic data analysis was performed in R IDE using a custom algorithm (https://github.com/asya-minervina/HLA) which aligns raw sequencing reads to HLA sequence database (extracted from IMGT/HLA database), get rid of PCR and sequencing errors and assemble HLA genotype of the donor.

### Sample collection for scRNA-Seq

For single-cell transcriptome analysis, we selected two donors in whom a substantial proportion of NK cells expressed the activating receptor KIR2DS4. The freshly isolated NK cells were stained with antibodies CD56-PE-Vio770, KIR2DS1-AF488, KIR2DS4-Vio-Blue, KIR2DL1-APC, NKG2C-FITC or NKG2C-PE, KIR3DL2-PE, CD3-PE, CD14-PE, and CD20-PE (Table 3), and using a FACSVantage DiVa cell sorter equipped with lasers: 405, 488, 590, and 633 nm (Becton Dickinson, USA). From donor 1 (D01), KIR2DS4^+^NKG2C^−^ and KIR2DS4^+^NKG2C^+^ NK cells (KIR3DL2^−^CD3^−^CD14^−^CD20^−^CD56^+^) were sorted, while from Donor 2 (D02) KIR2DS4^+^KIR2DS1^−^KIR2DL1^−^NKG2C^−^ and KIR2DS4-KIR2DL1C^245+^ NK cells (CD3^−^CD14^−^CD20^−^CD56^+^) were sorted. Preparation of the single-cell libraries was done immediately after collection of the resulting samples.

### scRNA-Seq library preparation

Four scRNA-Seq libraries were prepared using the Chromium Next GEM Single Cell 5’ Reagent Kits v2 (10x Genomics, USA) following the manufacturer’s instructions and purified using AMPure XP beads (Beckman Coulter, USA). Library size and quality were analyzed using a Fragment Analyzer (Advanced Analytical, Ames, Iowa, USA) before fragmentation and after final purification. The adapter ligation was done using MGIEasy PCR-Free DNA Library Prep Set (Shenzhen MGI Biological Electronic Technology Co, China). Final library concentrations were measured on a Qubit 2.0 fluorometer (Thermo Fisher, USA). Libraries were sequenced on a BJI platform (BJI, China).

### scRNA-Seq data analysis

scRNA-Seq reads were mapped to the GRCh38 reference genome with cellranger count, generating gene-per-cell count tables as output for further analysis. The original reference included only KIRs from haplotype A. To extract the KIRs counts from the haplotype B we manually searched the necessary sequences and calculated the counts under each barcode. Cellrenger output scRNA-Seq counts were imported into Seurat5 and filtered for outliers regarding frequency of mitochondrial transcripts (<5–6%), total number of transcripts (<15,000) and number of genes (<5,000) per cell. Contaminating erythrocytes and B cells were excluded based on the expression of HBA1/HBA2 and IGJ, respectively. The manually calculated KIR counts were imported into Seurat5 object metadata. Originally cellranger calculated counts of KIRs were removed. Next, counts were normalized using LogNormalize and used for principal-component analysis (PCA) on the 2,000 most variable genes. The number of principal components used for UMAP embedding and clustering was chosen based on elbow plots for each individual. The resolution was equal to 1.5. The normalized, KIRs corrected RNA assay was used for all differential expression analyses.

### KIR2DS4 clonal culturing

KIR2DS4⁺ *ex vivo* NK cells were sorted using a FACSVantage DiVa cell sorter (BD BioSciences, USA). Clonal cultures of KIR2DS4⁺ NK cells were generated by “single-cell” mode of sorting, with one cell deposited per well of a 96-well plate. To stimulate proliferation and activation of KIR2DS4⁺ NK cells, irradiated K562 feeder cells expressing membrane-bound IL-21 (K562-mbIL21) were used in combination with IL-2 (Hoffmann La-Roche, Switzerland) at 100 units/mL. Cells were cultured in the complete medium described above. After three to four weeks, phenotypic analysis of the expanded clones was performed. Surface expression of the following markers was assessed: KIR2DS4, KIR2DL1, NKG2C, NKG2A, and CD57. Based on phenotyping results and cell counts, clones with sufficient cell numbers were collected in triplicate into RLT buffer (Qiagen) for subsequent library preparation and RNA sequencing to characterize transcriptional differences between KIR2DS4⁺ clonal cultures.

### RNA Preparation and Sequencing of KIR2DS4^+^ NK cell clones

Total RNA was extracted from the samples with TRIzol (Invitrogen, Carlsbad, USA). The following conversion of total RNA into a cDNA was performed using mRNA-Seq Lib Prep Module for Illumina (ABclonal, China) according to the manufacturer’s instructions. Adaptor ligation and the addition of indexes were conducted using RNA Adapter Module 96 Index for Illumina (ABclonal, China). The final average fragment size was 300 bases. Each library was then subjected to paired-end sequencing on the Illumina NovaSeq 6000.

### Proliferation Test

Freshly isolated CD3-depleted PBMCs were stained with 5 μM CFSE (Molecular Probes, Eugene, Oregon, USA) in 100 μL of PBS (PanEco, Russia) and subsequently incubated for 15 min at 37°C protected from light. NK cells were washed three times in RPMI-1640 (PanEco, Russia) supplemented with 10% FBS (Hyclone, Utah, USA). CFSE^high^ NK cells were cultured in three replicates for each donor for 7 days in a complete medium. The medium was changed once on Day 4. The control samples were cultured without any stimulation. SARS-CoV-2 or EBV peptides (Table 2) were added on Days 0 and 4 at 100 μM concentration. To exclude the dead cells during analysis, the Sytox-VioBlue dye (Invitrogen, USA) was used. Highly proliferating CFSE^low^CD56^+^CD3^−^CD14^−^CD20^−^Sytox^−^NK cells (CFSE^low^ NK cells) were isolated on day 7 by cell sorting using a FACSVantage DiVa cell sorter. CFSE^low^ NK cells of donors were collected in three replicates in the RLT lysis buffer (QIAGEN, Germany) for RNA sequencing.

### RNA Preparation and Sequencing of CFSE^low^ NK cells

Total RNA was extracted from the samples with TRIzol (Invitrogen, Carlsbad, USA). Treatment with DNase was performed to avoid interference with genomic DNA (Thermo Fisher Scientific, MA, USA). The following conversion of total RNA into a cDNA was performed using SMART-Seq® v4 Ultra® Low Input RNA Kit for Sequencing (Takara Bio, USA) according to the manufacturer’s instructions. Fragmentation, adaptor ligation and the addition of indexes were conducted using Nextera XT DNA Library Prep Kit (Illumina, USA). The final average fragment size was 300 bases. Each library was then subjected to paired-end sequencing on the Illumina NovaSeq 6000.

### Transcriptomic Analysis

Raw RNA-sequencing data quality was assessed by FASTQC. Paired-end RNA-sequencing reads were aligned to version 38 of the Ensembl annotation of the human reference genome with the following unique gene hit counts calculation using Salmon. DESeq2 was used to analyze the data and perform the differential expression assay. Genes with absolute log2 fold change (LFC) > 0.58 and Padj < 0.05 were considered as DEGs for each comparison. For transcriptomic analysis of proliferating NK cells additional filtering basemean > 40 was applied.

### In silico modeling

The sequence and 3D crystal structure of HLA-KIR complexes were obtained from the PDB. Prediction of the structure of complexes KIR2DS4-CRA-HLA-C*06 and addition of the glycan structures were conducted using AlphaFold server. The obtained models were arranged by the predicted local distance difference test (pLDDT), and the model with highest confidence was selected for further visualization. Visualization of obtained structures was performed in the PyMol molecular visualization system (Version 3.0.3, Schrödinger, LLC). The polar contacts were identified by the “find” function and the distances were highlighted in the figure. Due to the variability in KIR2DS4 proteins, which may have the different length of the transmembrane part, the aminoacids numbering was started with the first amino acid in the crystal structure.

### Statistical analysis

Data were analyzed using Microsoft Excel, FlowJo (FlowJo X, Oregon, USA), and GraphPad Prism (version 8.4.3, GraphPadSoftware, USA) and R language. The normality of the data was assessed by the Shapiro-Wilk test. Figures represent mean ± standard deviation; statistical analysis using one-way-ANOVA for multiple comparisons were performed. p value < 0.05 was considered statistically significant. The correlation assay was performed by identification of Pearson coefficients.

## Supplementary Figures

**Supplementary Figure 1. scRNA-Seq of sorted KIR2DS4^+^ NK cells of Donor 2.**

**Supplementary Figure 2. scRNA-Seq of sorted KIR2DS4^+^NKG2C^+^ NK cells of Donor 1.**

**Supplementary Figure 3. scRNA-Seq of sorted KIR2DS4^+^NKG2C^−^ NK cells of Donor 1.**

**Supplementary Figure 4. KIR2DS4^+^ NK cells characteristics.**

## Author contributions

DMC conceived and designed the study. MOU performed and analyzed the data for most of the experiments, with support from DMC, IAS, MAS, EIK, EN, EK. MOU and DMC performed the bioinformatics analyses. MAS performed HLA-genotyping. MOU prepared the figures. DMC, EIK, OVB, YV provided key resources. DMC, EIK, OVB supervised the work. DMC and MOU wrote the paper with important input from EIK. All authors read and approved the manuscript.

## Data availability

Data used in the analysis will be available in figshare under DOI: 10.6084/m9.figshare.32394207.

