## Supplemental for "Towards understanding of NK cell antigenic specificity"

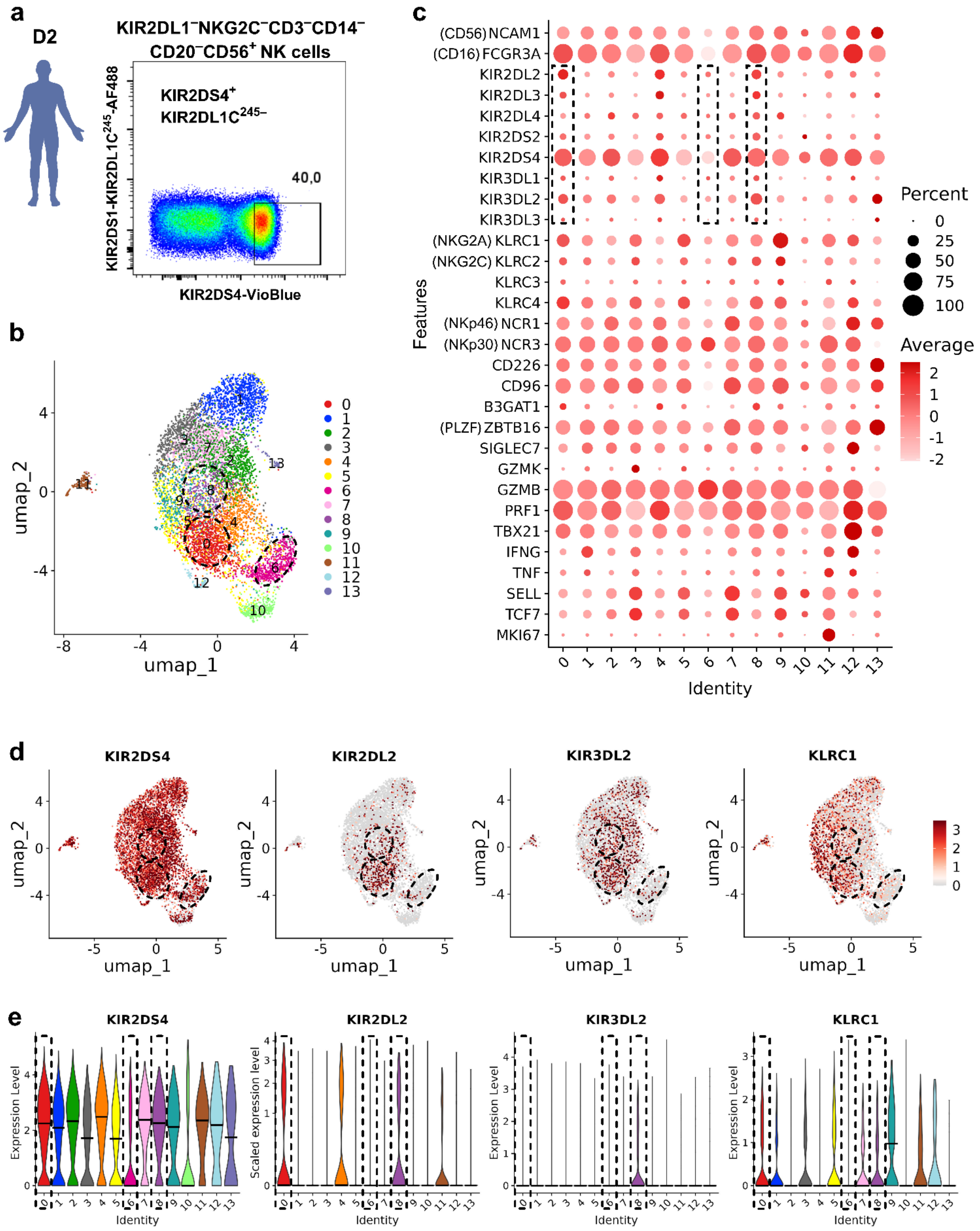


**Supplementary Figure 1. scRNA-Seq of sorted KIR2DS4⁺ NK cells of Donor 2. a.** Sorting gates to obtain KIR2DS4⁺ NK cells. **b**. UMAP at high resolution. **c**. Dotplot with NK cell receptor genes and genes for general characterization. **d**. UMAP with selected NK cell receptor genes. **e**. Violin plots with the same genes.

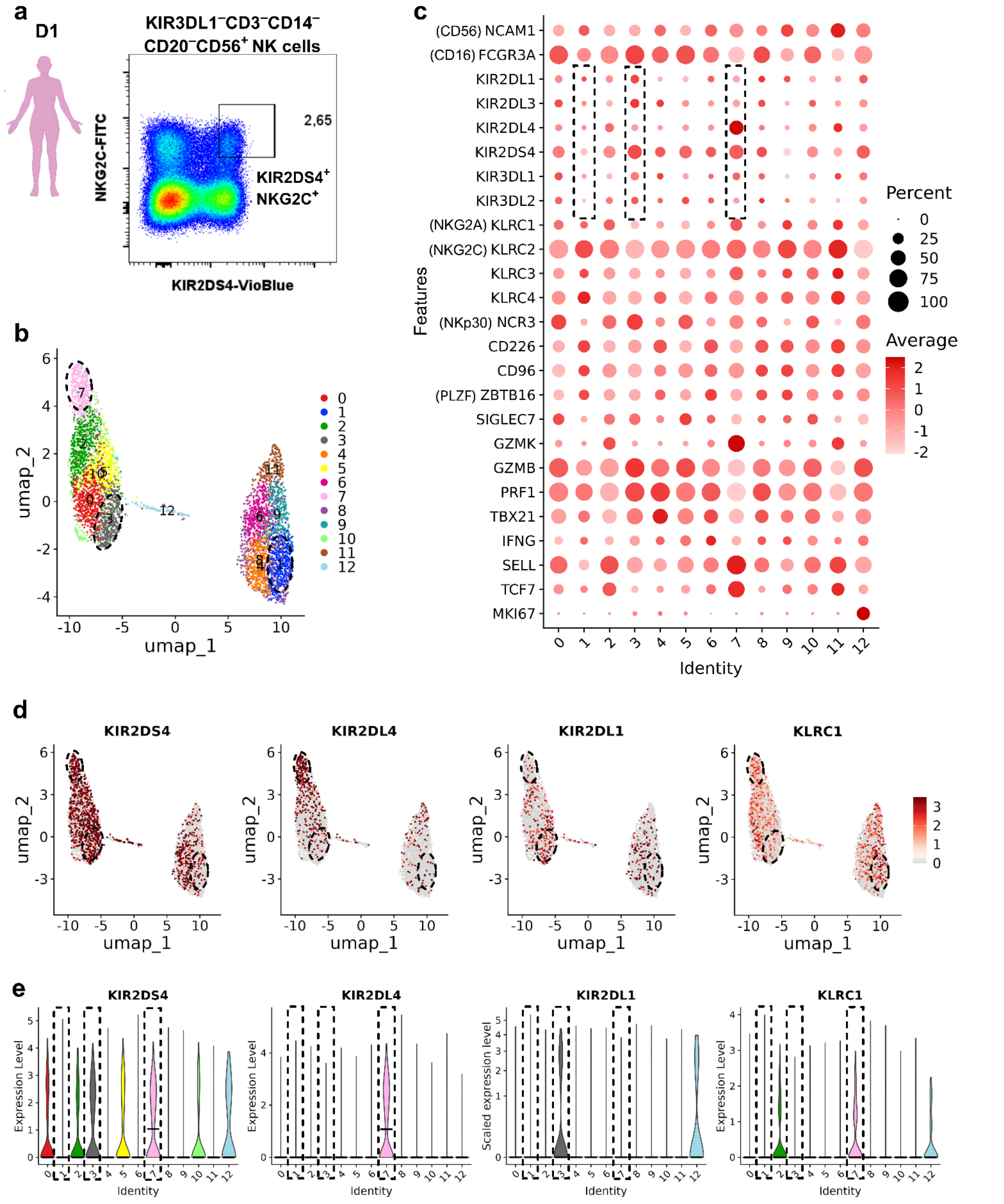


**Supplementary Figure 2. scRNA-Seq of sorted KIR2DS4⁺NKG2C⁺ NK cells of Donor 1. a.** Sorting gates to obtain KIR2DS4⁺NKG2C⁺ NK cells. **b**. UMAP at high resolution. **c**. Dotplot with NK cell receptor genes and genes for general characterization. **d**. UMAP with selected NK cell receptor genes. **e**. Violin plots with the same genes.

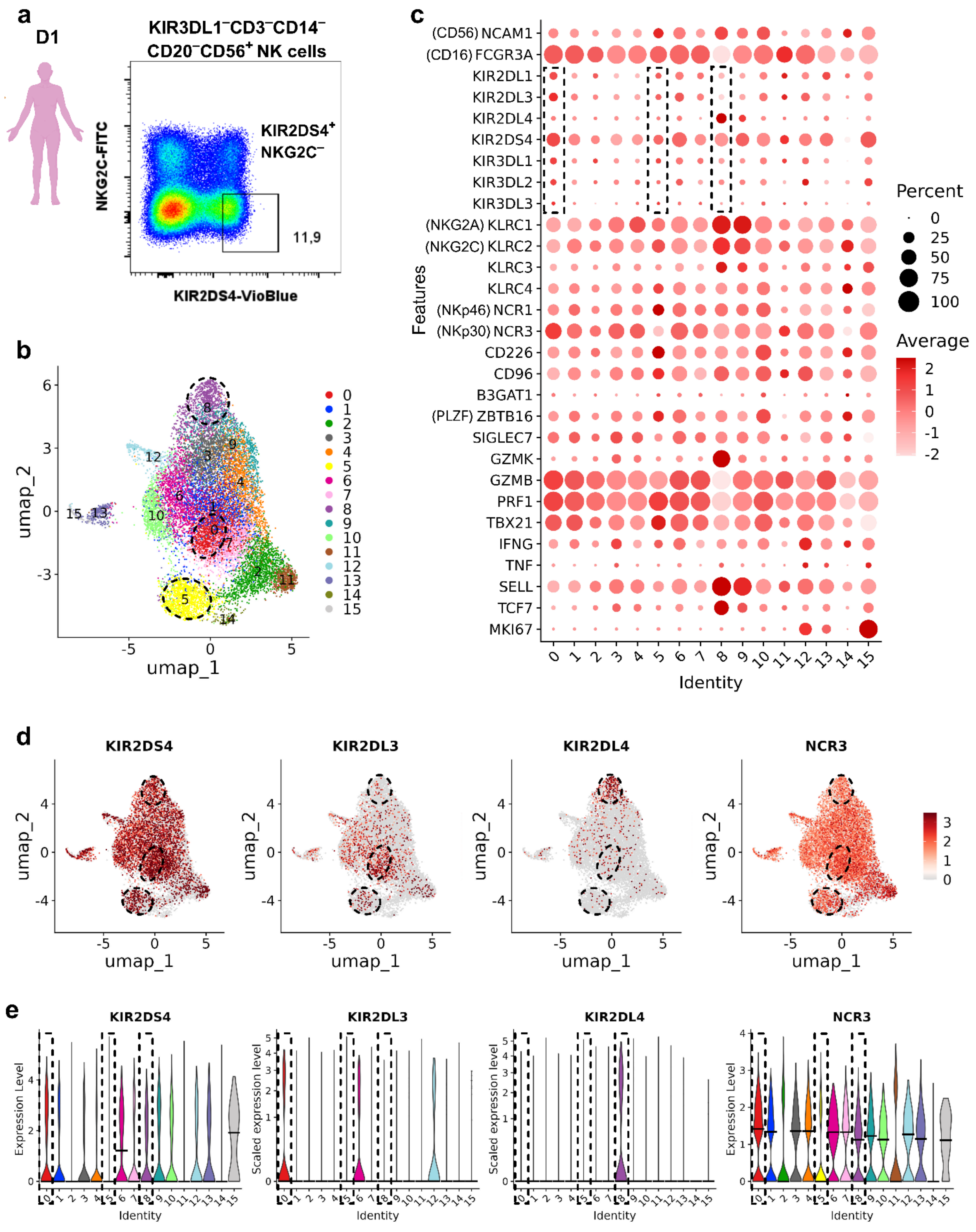


**Supplementary Figure 3. scRNA-Seq of sorted KIR2DS4⁺NKG2C⁻ NK cells of Donor 1. a.** Sorting gates to obtain KIR2DS4⁺NKG2C⁻ NK cells. **b**. UMAP at high resolution. **c**. Dotplot with NK cell receptor genes and genes for general characterization. **d**. UMAP with selected NK cell receptor genes. **e**. Violin plots with the same genes.

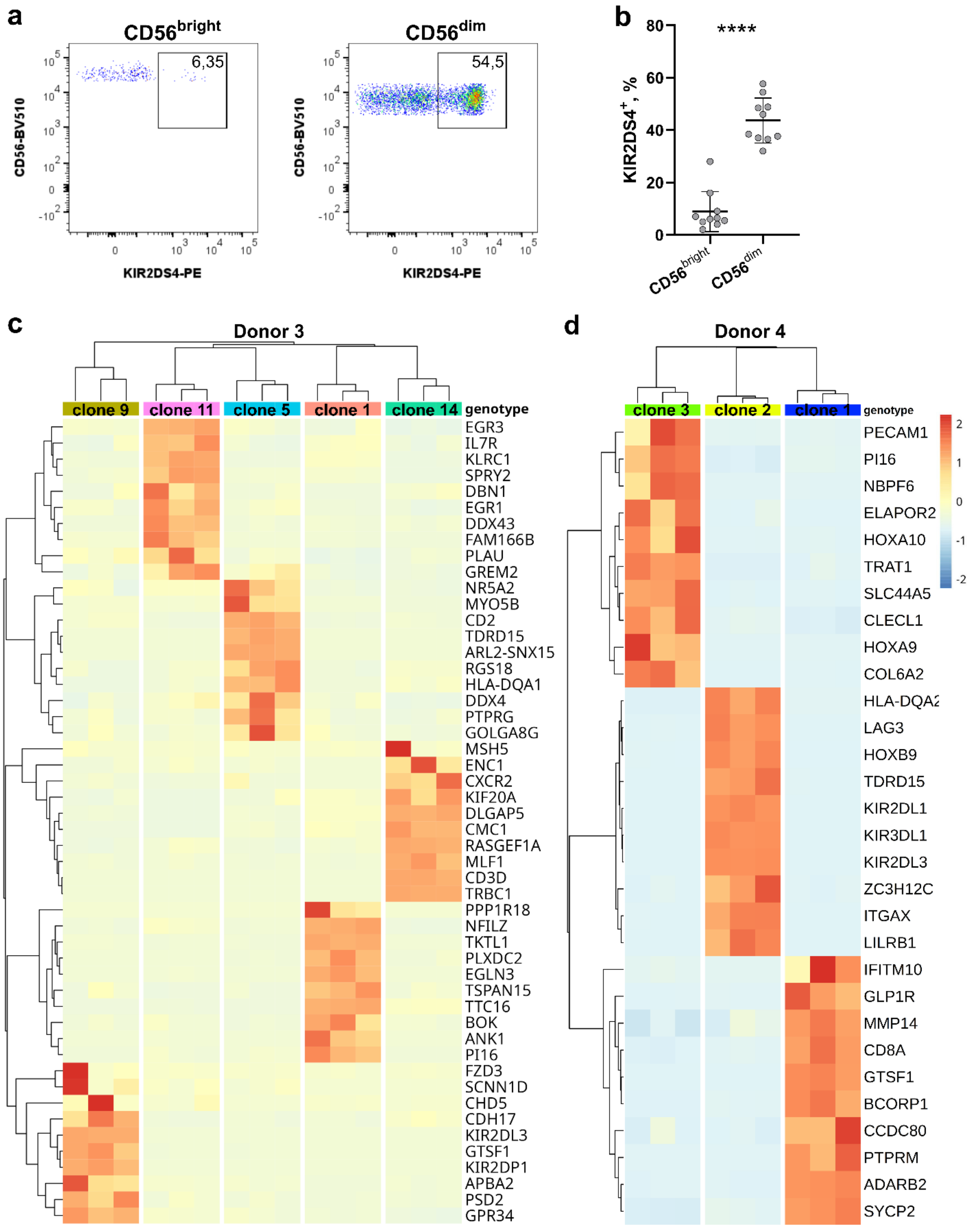


**Supplementary Figure 4. KIR2DS4$^{+}$ NK cells characteristics. a**. Typical flow cytometry distribution of KIR2DS4$^{+}$ in CD56$^{bright}$ and CD56$^{dim}$ NK cells. **b**. Proportions of KIR2DS4$^{+}$ NK cells in CD56$^{bright}$ and CD56$^{dim}$ NK cells in 10 donors participated in the study. **c**,**d**. Heatmap of top 10 differentially expressed genes in each clone for Donor 3 (c) and Donor 4 (d).
